# Aggregation of CAT tails blocks their degradation and causes proteotoxicity in *S. cerevisiae*

**DOI:** 10.1101/687319

**Authors:** Cole S. Sitron, Joseph H. Park, Jenna M. Giafaglione, Onn Brandman

**Author notes:** These authors contributed equally to the work (Cole S. Sitron, Joseph H. Park).

## Abstract

The Ribosome-associated Quality Control (RQC) pathway co-translationally marks incomplete polypeptides from stalled translation with two signals that trigger their proteasome-mediated degradation. The E3 ligase Ltn1 adds ubiquitin and Rqc2 directs the large ribosomal subunit to append carboxy-terminal alanine and threonine residues (CAT tails). When excessive amounts of incomplete polypeptides evade Ltn1, CAT-tailed proteins accumulate and can self-associate into aggregates. CAT tail aggregation has been hypothesized to either protect cells by sequestering potentially toxic incomplete polypeptides or harm cells by disrupting protein homeostasis. To distinguish between these possibilities, we modulated CAT tail aggregation in *Saccharomyces cerevisiae* with genetic and chemical tools to analyze CAT tails in aggregated and un-aggregated states. We found that enhancing CAT tail aggregation induces proteotoxic stress and antagonizes degradation of CAT-tailed proteins, while inhibiting aggregation reverses these effects. Our findings suggest that CAT tail aggregation harms RQC-compromised cells and that preventing aggregation can mitigate this toxicity.

## Introduction

Failed rounds of translation produce incomplete, potentially toxic polypeptides that organisms across all clades of life have evolved responses to degrade (Brandman and Hegde, 2016; Defenouillère and Fromont-Racine, 2017; Ikeuchi et al., 2018; Joazeiro, 2019; Moore and Sauer, 2007). In prokaryotes, the primary degradative response involves a tRNA-mRNA hybrid molecule (tmRNA) (Moore and Sauer, 2007). The tmRNA enters stalled ribosomes, re-initiates translation elongation with its tRNA moiety and switches the ribosome’s template to its mRNA moiety (Moore and Sauer, 2007). This prompts the ribosome to synthesize a tmRNA-encoded tag on the incomplete polypeptide’s C-terminus that marks it for proteolysis (Moore and Sauer, 2007). The eukaryotic response, called <u>R</u>ibosome-associated <u>Q</u>uality <u>C</u>ontrol (RQC), begins when a set of factors recognize ribosomes that have stalled on the same mRNA and collided into each other (Ikeuchi et al., 2019; Juszkiewicz et al., 2018; Simms et al., 2017). These factors then split the ribosomes into their large and small subunits, leaving the incomplete polypeptide (RQC substrate) tethered to the large subunit (Juszkiewicz and Hegde, 2017; Lyumkis et al., 2014; Matsuo et al., 2017; Pisareva et al., 2011; Shao et al., 2013; Shoemaker et al., 2010; Sitron et al., 2017; Sundaramoorthy et al., 2017; Tsuboi et al., 2012). The E3 ligase Ltn1 binds to the large subunit and ubiquitylates the incomplete polypeptide, marking it for proteasomal degradation (Bengtson and Joazeiro, 2010; Brandman et al., 2012; Defenouillère et al., 2013; Shao et al., 2013; Verma et al., 2013).

Disruption of tmRNA or Ltn1 compromises the cell’s ability to degrade incomplete polypeptides and reduces survival under stresses that increase translational stalling (Abo et al., 2002; Choe et al., 2016; Defenouillère et al., 2013; Kostova et al., 2017; Lytvynenko et al., 2019; Moore and Sauer, 2005; Oussenko et al., 2005). This deficit in fitness at the cellular level has clinically-relevant consequences. tmRNA deficiency prevents growth of some disease-causing prokaryotes (e.g. *Mycoplasma genitalium* and *Staphylococcus aureus*) and decreases the virulence of others (e.g. *Salmonella enterica* and *Streptococcus pneumoniae*) (Chaudhuri et al., 2009; Glass et al., 2006; Julio et al., 2000; Mann et al., 2012). In metazoan eukaryotes, mutations to *LTN1* or perturbations that introduce large influxes of RQC substrates lead to neurodegeneration (Chu et al., 2009; Ishimura et al., 2014; Wu Z, Tantray I, Lim J, Chen S, Li Y, Davis Z, Sitron C, Dong J, Gispert S, Auburger G, Brandman O, Bi X, Snyder M, Lu B., 2019). Each of these phenotypes highlights the central role that tmRNA and Ltn1 play in maintaining protein homeostasis and avoiding the toxicity associated with compromised co-translational quality control.

A conserved back-up degradation pathway mediated by Rqc2 and its prokaryotic homologs mitigates some of the toxicity associated with loss of tmRNA or Ltn1 function (Lytvynenko et al., 2019; Sitron and Brandman, 2019). Rqc2 homologs bind the large ribosomal subunit and direct it to elongate the incomplete polypeptide’s C-terminus with either alanine (“Ala tails” in prokaryotes) or both alanine and threonine residues (“CAT tails” in yeast) (Lytvynenko et al., 2019; Osuna et al., 2017; Shen et al., 2015). Metazoan CAT tails may include a more diverse repertoire of amino acids (Wu Z, Tantray I, Lim J, Chen S, Li Y, Davis Z, Sitron C, Dong J, Gispert S, Auburger G, Brandman O, Bi X, Snyder M, Lu B., 2019). These extensions, made without a small subunit or mRNA, act as “degrons” to mark incomplete polypeptides for degradation by the bacterial protease ClpXP or the eukaryotic proteasome (Lytvynenko et al., 2019; Sitron and Brandman, 2019).

Loss of Ltn1 function results in a buildup of CAT-tailed (“CATylated”) proteins (Shen et al., 2015). The CAT tails on these proteins can homo-typically associate and form aggregates of CATylated proteins, which have been observed in yeast and metazoans (Choe et al., 2016; Defenouillère et al., 2016; Wu Z, Tantray I, Lim J, Chen S, Li Y, Davis Z, Sitron C, Dong J, Gispert S, Auburger G, Brandman O, Bi X, Snyder M, Lu B., 2019; Yonashiro et al., 2016). The presence of threonine in yeast CAT tails increases their aggregation propensity (Yonashiro et al., 2016). It is thereby an attractive hypothesis that CAT tail aggregation has an adaptive function during stress. For instance, aggregation may protect the proteome by sequestering potentially harmful undegraded RQC substrates when Ltn1 becomes overburdened. Alternatively, it has been hypothesized that CAT tail aggregates disrupt cellular fitness by depleting the pool of chaperones (Choe et al., 2016; Defenouillère et al., 2016; Yonashiro et al., 2016), and thereby contribute to the proteotoxicity associated with Ltn1 failure. No study has evaluated these hypotheses by controlling the aggregation state of CAT tails and testing whether CAT tail aggregation plays a beneficial or harmful role during stress.

We used chemical and genetic approaches in *Saccharomyces cerevisiae* to modulate CAT tail aggregation propensity in Ltn1-deficient cells and determine how aggregation affects: 1) the function of CAT tails as degrons and 2) proteotoxic stress. We found that under conditions favoring CAT tail aggregation (elevated Rqc2 levels), CAT tails lose their degron function and induce proteotoxic stress. Conversely, we observed that increasing CAT tail solubility (with guanidinium hydrochloride or by genetic perturbation of RNA Polymerase III function) restores CAT tail-mediated degradation and reduces proteotoxic stress. Our work demonstrates that CAT tail aggregation contributes to the toxicity of Ltn1 disruption rather than aiding in adaptation to stress.

## Results

### CAT tail aggregation is associated with compromised CAT tail degradation

We began our investigation into the effects of aggregation on CAT tail-mediated degradation by validating the model RQC substrate RQCsub. RQCsub consists of green fluorescent protein (GFP) attached via an inert linker including a tobacco etch virus (TEV) protease site to twelve stall-inducing arginine CGN codons (Figure 1A) (Letzring et al., 2010). Translation of RQCsub produces a stalled GFP-linker-arginine “arrest product” (Brandman et al., 2012). The RQCsub arrest product accumulated at low levels in wt strains by SDS-PAGE (Figure 1A). The arrest product increased in abundance after we compromised RQC function by deleting either of two RQC genes: the CAT tail-elongating factor *RQC2* or the E3 ubiquitin ligase *LTN1* (Figure 1A). This result indicates that Rqc2 and Ltn1 contribute to RQCsub degradation. While deletion of *RQC2* or *LTN1* stabilized RQCsub, the resultant protein products displayed different mobilities: RQCsub from *rqc2*Δ cells migrated as a crisp band and that from *ltn1*Δ cells migrated as a higher molecular weight smear (Figure 1A). This smear collapsed into a single band after additional deletion of *RQC2* in the *ltn1*Δ strain (Figure 1A), consistent with the smear containing CATylated RQCsub. Taken together, these results confirm that RQCsub is degraded via RQC and CATylated by Rqc2.

**Figure 1.**
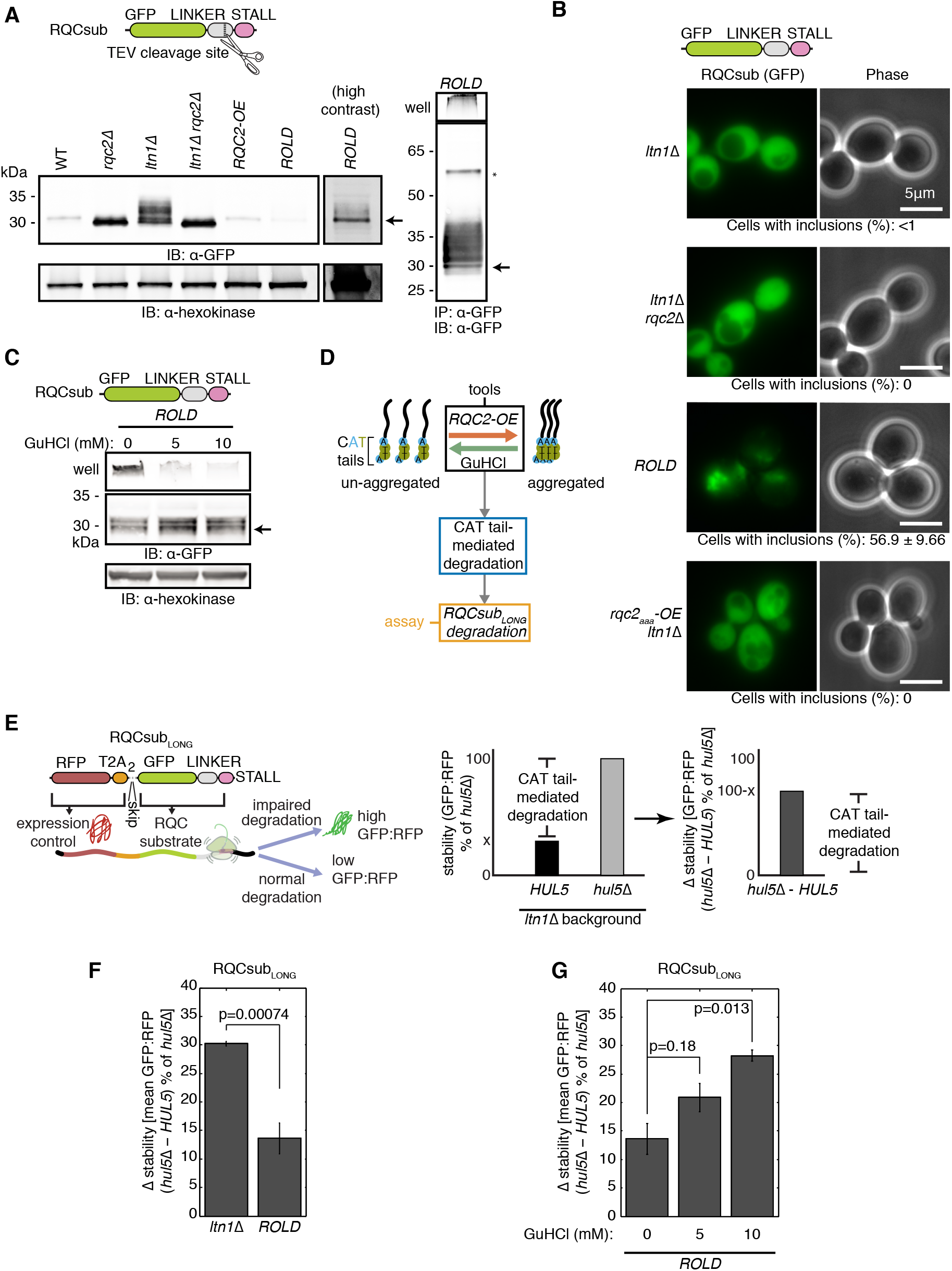
Aggregation compromises CAT tail degradation. (**A**) Left, whole cell immunoblots (IBs) of lysates containing the model RQC substrate RQCsub (schematic above). Right, IB of RQCsub immunoprecipitated (IPed) from *RQC2* overexpression and *LTN1* deletion (*ROLD*) lysate. Arrow indicates molecular weight of RQCsub without CAT tails. Asterisk denotes the full-length RQCsub protein product, produced when ribosomes translate through the stall sequence (region past the stall not pictured in schematic). GFP, green fluorescent protein. OE, overexpression. (**B**) Fluorescence microscopy of cells expressing RQCsub with percentages of cells containing observable GFP inclusions reported below. (**C**) IB of RQCsub expressed in *ROLD* cultures grown in guanidinium hydrochloride (GuHCl). Arrows as in **A**. (**D**) Schematic of experimental approach for measuring the effect of aggregation of CAT tail-mediated degradation. (**E**) Left, schematic of expression-controlled model RQC substrate RQCsub_LONG_. Center, definitions of “stability,” “Δ stability,” and “CAT tail-mediated degradation.” (**F**) and (**G**) Δ stability measurements of RQCsub_LONG_ expressed in *ltn1*Δ and *ROLD* cells after *HUL5* deletion to assess CAT tail-mediated degradation. P-values above bars are derived from t-tests for particular contrast, measuring how significantly different *HUL5* deletion-induced stabilization is between either *ltn1*Δ and *ROLD* backgrounds in F or between different GuHCl concentrations in **G**. Error bars represent the standard error of the mean (s.e.m.).

To vary the aggregation propensity of CAT tails, we overexpressed *RQC2*, which has previously been shown to increase CAT tail aggregation (Yonashiro et al., 2016). Overexpression of *RQC2* with the *TDH3* promoter increased Rqc2 levels 28.4-fold relative to the native promoter, as assessed by levels of red fluorescent protein (RFP)-tagged Rqc2 (Figure 1-figure supplement 1A). In the *<u>R</u>QC2* <u>o</u>verexpression and *<u>L</u>TN1* <u>d</u>eletion strain (hereafter named “*ROLD*”), a proportion of RQCsub remained in the well rather than migrating into an SDS-PAGE gel (Figure 1A) despite prior boiling in dodecylsulfate detergent. This detergent-resistant species was more prominent in *ROLD* than in *ltn1*Δ lysates (Figure 1-figure supplement 1B). TEV treatment, which cleaves RQCsub’s C-terminus to remove its CAT tail, enabled RQCsub purified from *ROLD* cells to migrate as a single band into the gel (Figure 1-figure supplement 1C). The properties of this detergent-resistant species are consistent with *RQC2* overexpression reducing the solubility of CATylated RQCsub in the *ltn1*Δ background. Microscopy revealed that RQCsub had a diffuse localization in the majority of *ltn1*Δ cells (<1% of cells with inclusions) but became more punctate in *ROLD* cells (56.9% of cells with inclusions) (Figure 1B). To test whether these inclusions depended on CATylation, we disrupted CATylation by deleting *RQC2* or overexpressing a CATylation-incompetent *rqc2_aaa_*allele (Shen et al., 2015) instead of *RQC2-WT*. Neither of these perturbations (*rqc2*Δ *ltn1*Δ nor *rqc2_aaa_-OE ltn1*Δ) induced the punctate RQCsub localization we observed in *ROLD* cells (Figure 1B), confirming that formation of RQCsub inclusions requires CATylation. These results demonstrate that the *ROLD* genetic background can be used to enhance CAT tail aggregation.

We next sought a tool to inhibit CAT tail aggregation. Millimolar concentrations of the chaotrope guanidinium hydrochloride (GuHCl) in yeast growth medium solubilize CAT tail aggregates and prevent propagation of aggregated yeast prions (Byrne et al., 2007; Defenouillère et al., 2016; Eaglestone et al., 2000; Ness et al., 2002). When we treated *ROLD* cells with GuHCl, we observed that RQCsub still localized to inclusions by microscopy (Figure 1-figure supplement 1D). However, monitoring RQCsub mobility in SDS-PAGE assessed how GuHCl affected RQCsub more sensitively than microscopy, revealing that GuHCl treatment enabled a larger proportion of RQCsub extracted from *ROLD* cells to migrate into the resolving gel rather than remaining in the well (Figure 1C). GuHCl thus reduces the detergent insolubility of CAT tails and can thereby serve as a tool to inhibit CAT tail aggregation.

Single-color model RQC substrates like RQCsub, while useful for visualizing CAT tails by SDS-PAGE and microscopy, have limited utility in degradation measurements due to noisy expression (Sitron and Brandman, 2019). To assess how aggregation affects degradation of CATylated RQC substrates, we utilized the two-color model RQC substrate RQCsub_LONG_ (Sitron and Brandman, 2019) (Figure 1D,E). RQCsub_LONG_ includes an internal expression control, RFP followed by viral T2A peptides, at the N-terminus of GFP-linker-arginine (Figure 1E). Ribosomes translating RQCsub_LONG_ produce RFP then skip formation of a peptide bond within the T2A sequence (Donnelly et al., 2001; Szymczak and Vignali, 2005), severing RFP from the GFP-linker-arginine RQC substrate (Figure 1E). As a result, each round of translation produces RFP and GFP stoichiometrically, but only GFP becomes an RQC substrate (Figure 1E). When we analyzed GFP and RFP separately, we observed that each signal was lower in *ROLD* than in *ltn1*Δ cells (Figure 1-figure supplement 2A). This loss of fluorescent signal was especially evident for the expression control RFP (Figure 1-figure supplement 2A), indicative of reduced RQCsub_LONG_ translation in the *ROLD* background. This reduction in translation may explain why we observed lower RQCsub levels in the *ROLD* background compared to *ltn1*Δ by SDS-PAGE (Figure 1A) and highlights the need for a precise way to measure degradation that is not confounded by changes in translation.

We then controlled for these differences in translation to quantify how increased CAT tail aggregation (in the *ROLD* strain) affects degradation of CATylated RQCsub_LONG_. The ratio of GFP:RFP at the single-cell level serves as an internal, expression-controlled readout of RQCsub_LONG_ stability (Figure 1E). CAT tail-mediated degradation of RQCsub_LONG_ in the absence of Ltn1 requires the proteasome-associated E3 ubiquitin ligase Hul5 and the proteasome (Sitron and Brandman, 2019). Therefore, comparing stability before and after *HUL5* deletion quantifies CAT tail-mediated degradation (Figure 1E). RQCsub_LONG_ stability did not change after treatment of *ltn1*Δ *hul5*Δ or *ROLD hul5*Δ cells with the proteasome inhibitor bortezomib (Figure 1-figure supplement 2B), supporting the requirement for Hul5 in CAT tail-mediated degradation of RQCsub_LONG_ by the proteasome. *HUL5* deletion stabilized RQCsub_LONG_ by 30.2% in *ltn1*Δ and 13.6% in *ROLD* backgrounds (Figure 1F), consistent with *RQC2* overexpression reducing CAT tail-mediated RQCsub_LONG_ degradation. These results demonstrate that the *ROLD* genetic background, which potentiates CAT tail aggregation, impairs CAT tail-mediated degradation.

To ensure that it is aggregation (and not another effect of *ROLD*) that reduces CAT tail-mediated degradation, we inhibited aggregation with GuHCl and measured RQCsub_LONG_ stability. We observed that GuHCl treatment had two effects on RQCsub_LONG_ degradation in *ROLD* strains. First, increasing concentrations of GuHCl increased the magnitude of the effect of *HUL5* deletion on the stability of RQCsub_LONG_ (from 13.6% to 20.9% to 28.2% at 0, 5, and 10 mM GuHCl respectively; Figure 1G), suggesting that treating *ROLD* cells with GuHCl rescues degradation of RQCsub_LONG_. Second, the magnitude of bortezomib-induced RQCsub_LONG_ stabilization increased with increasing GuHCl when Hul5 was intact (Figure 1-figure supplement 2C). This result indicates that the degradation that GuHCl rescued involved both Hul5 and the proteasome. Taken together, these data imply that the aggregation-conducive *ROLD* background impairs CAT tail-mediated degradation, and solubilization of CAT tails rescues this impairment. We thereby posit that CAT tail aggregation antagonizes CAT tail-mediated degradation.

### CAT tails exert proteotoxic stress in their aggregated state

To determine how aggregation affects the proteotoxicity of CAT tails, we measured proteotoxic stress after varying CAT tail aggregation propensity (Figure 2A). Proteotoxic stress activates the transcription factor Heat Shock Factor 1 (Hsf1), which then orchestrates transcription of chaperones and other proteostasis-restoring factors (Åkerfelt et al., 2007; Morimoto, 2011). Hsf1 activation scales with the magnitude of proteotoxic stress and a previously-published reporter can measure the degree of activation (Brandman et al., 2012). This reporter consists of GFP under the control of an artificial promoter containing four heat shock elements (Sorger and Pelham, 1987), the consensus binding site for Hsf1 (Figure 2B). *LTN1* deletion alone activated the Hsf1 reporter 2.07-fold relative to wt, while additional *RQC2* overexpression (*ROLD*) led to 22.6-fold activation (Figure 2B). The potent activation observed in the *ROLD* background required *LTN1* deletion, as *RQC2* overexpression in the wt background weakly affected the reporter (Figure 2B). Thus, the combination of *RQC2* overexpression and *LTN1* deletion, which potentiates CAT tail aggregation (Figure 1-figure supplement 1B) (Yonashiro et al., 2016), generates a synthetic interaction that results in elevated Hsf1 activity. This increase in Hsf1 activity is consistent with an increase in proteotoxic stress.

**Figure 2.**
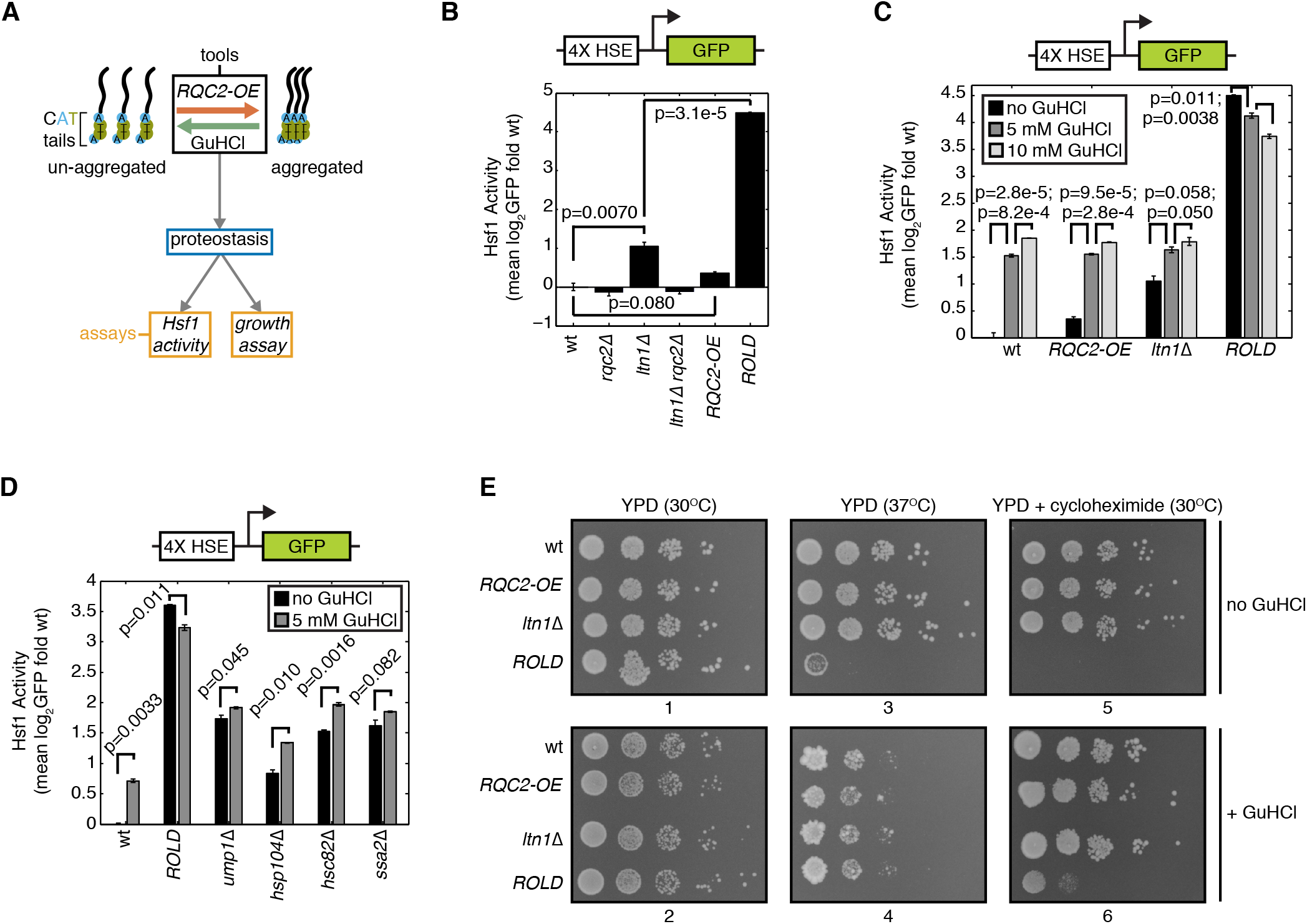
Aggregated CAT tails are proteotoxic. (**A**) Schematic of experimental approach for assessing how aggregation influences the effect of CAT tails on proteostasis. (**B**), (**C**), and (**D**) Flow cytometry of cells containing an integrated reporter for Hsf1 activation (schematic above). Error bars represent s.e.m. from three independent cultures. P-values from indicated paired t-tests are indicated by lines between bars. HSE, heat shock element. (**E**) Spot assay of strains grown at 30°C, 37°C, and in 50 ng/ml cycloheximide with and without the presence of 5 mM GuHCl. Plate numbers indicated below images.

If CAT tail aggregation causes proteotoxicity, we reasoned that solubilization of CATylated proteins by GuHCl treatment might reduce this toxicity. GuHCl treatment activated the Hsf1 reporter in a dose-dependent manner in wt, *RQC2-OE*, and *ltn1*Δ strains (Figure 2C), indicating that GuHCl alone exerts some proteotoxic stress in cells. Despite this general activation, GuHCl decreased Hsf1 reporter activation in *ROLD* cells (Figure 2C), suggesting that solubilization of CAT tails alleviates the Hsf1 activation associated with the *ROLD* background. We tested the specificity of this response by measuring the effect of GuHCl on the Hsf1 reporter in other genotypes with high Hsf1 activity (Brandman et al., 2012) (Figure 2D). Hsf1 reporter deactivation was unique to the *ROLD* condition, as GuHCl increased Hsf1 reporter activation in *ump1*Δ, *hsp104*Δ, and *hsc82*Δ strains and had no significant effect in the *ssa2*Δ strain (Figure 2D). These results suggest that CAT tails exert proteotoxic stress more potently in their aggregated than in their soluble state.

To ensure that our Hsf1 measurements reflect perturbations to proteostasis, we analyzed how CAT tail aggregation affects cells’ sensitivity to stressors. At 30°C, all of the strains we examined (wt, *RQC2* overexpression, *ltn1*Δ, and *ROLD*) grew equally well in a spot assay in the absence and presence of 5 mM GuHCl (Figure 2E, plates 1 and 2). This result demonstrates that neither the *ROLD* background nor GuHCl impairs growth in the absence of stress. We then introduced two mild stressors to perturb proteostasis: the thermal stress of 37°C incubation or introduction of 50 ng/ml cycloheximide to increase translational stalling. Neither of these stresses induced a growth defect in the *RQC2* overexpression (*RQC2-OE*) or *ltn1*Δ strains relative to wt (Figure 2E, plates 3 and 5). By contrast, 37°C incubation and 50 ng/ml cycloheximide severely inhibited growth in the *ROLD* strain (Figure 2E, plates 3 and 5). Increased CAT tail aggregation thus co-occurs with sensitivity to thermal and ribosome stalling stresses. Additional GuHCl partially rescued growth of the *ROLD* strain in the presence of cycloheximide (Figure 2E, plate 5 versus plate 6). The combination of GuHCl and 37°C led to a synthetic growth defect in all strains (Figure 2E, plate 3 versus plate 4). Despite this growth defect, the *ROLD* strain grew comparably to wt, *RQC2-OE*, and *ltn1*Δ strains when exposed to both GuHCl and 37°C (Figure 2E, plate 4). Thus, CAT tail aggregation sensitizes cells to mild thermal and ribosome stalling stress. The agreement with our Hsf1 measurements (Figure 2B,C) suggests that CAT tail aggregation perturbs proteostasis under the conditions we measured rather than facilitating adaptation to stress.

### Disruption of CAT tail aggregation by Pol III perturbation restores CAT tail degradation and alleviates proteotoxicity

Given that GuHCl mildly reduced CAT tail aggregation, we sought an orthogonal perturbation that would more potently inhibit CAT tail aggregation. To find such a perturbation, we screened for mutations that reduced Hsf1 reporter activation in the RQC-compromised *rqc1*Δ *rps0a*Δ strain, which exhibits Hsf1 hyperactivation (Brandman et al., 2012). After crossing a genome-wide collection of hypomorphs into this background, we found that all hits other than *RQC2* (whose deletion blocks CATylation) were related to RNA Polymerase III (Pol III). These hits included a non-essential polymerase biogenesis factor, *BUD27* (Mirón-García et al., 2013), and essential Pol III subunits, *RPC17* and *RPC160*. Disruption of each of these by deletion or a “<u>d</u>ecreased <u>a</u>bundance by <u>m</u>RNA <u>p</u>erturbation” allele (DAmP) (Yan et al., 2008) nearly eliminated activation of the Hsf1 reporter in *ROLD* cells compared to wt (Figure 3-figure supplement 1A). To assess the specificity of this effect to RQC-related Hsf1 activation, we measured the effect of *BUD27* and *RPC17* perturbations in strains with elevated Hsf1 signaling (Figure 3A). Perturbations to *BUD27* and *RPC17* reduced Hsf1 reporter activity in wt, *ump1*Δ, *hsp104*Δ, *hsc82*Δ, and *ssa2*Δ strains (Figure 3A, black versus grey bars). However, the magnitude of this reduction in each strain was substantially less than the 20.7-fold and 13.7-fold reduction we observed in the *ROLD* background after disruption of *BUD27* and *RPC17*, respectively (Figure 3A). Furthermore, Hsf1 reporter activation increased in the Pol III-perturbed backgrounds upon deletion of *UMP1, HSP104, HSC82*, and *SSA2* relative to the Pol III-perturbed background strain (Figure 3-figure supplement 1B). Taken together, these data indicate that Pol III impairment reduces ROLD-induced Hsf1 activation without generally preventing Hsf1 activation in other genetic backgrounds.

**Figure 3.**
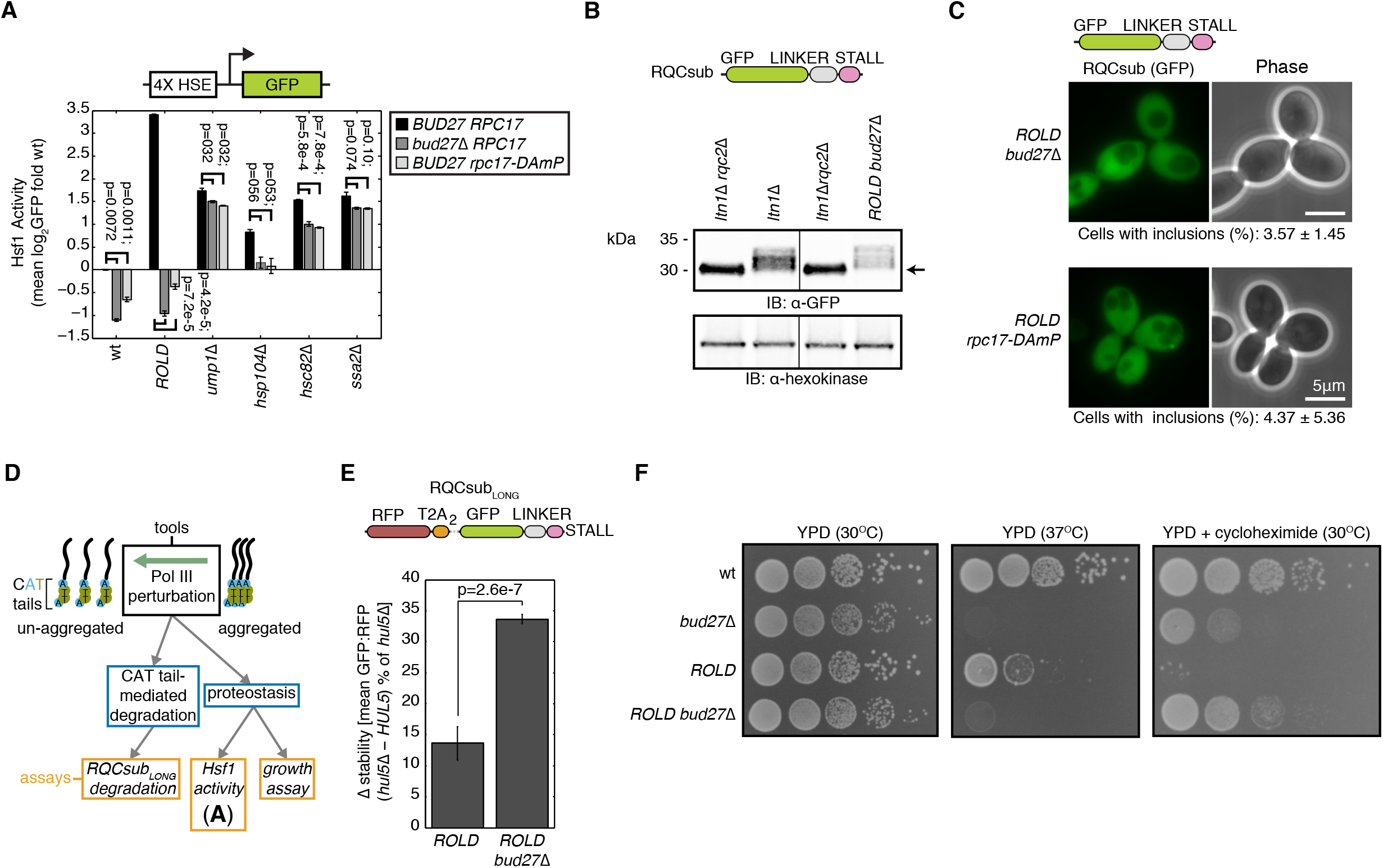
Pol III perturbation inhibits CAT tail aggregation, increases degron activity, and restores proteostasis. (**A**) Flow cytometry of cells containing an integrated reporter for Hsf1 activation. Error bars indicate s.e.m. from three independent cultures. P-values from paired t-tests indicated above bars. (**B**) IB of RQCsub expressed in indicated strains. (**C**) Fluorescence microscopy of cells expressing RQCsub. Percentages of cells containing GFP inclusions reported below micrographs. (**D**) Schematic of experimental strategy for measuring how disruption of aggregation modulates CAT tail-mediated degradation and CAT tails’ effect on proteostasis. (**E**) Δ stability measurements of RQCsub_LONG_ expressed in indicated strains after *HUL5* deletion to quantify CAT tail-mediated degradation. Error bars as in **A**. P-value indicates the result of a t-test for particular contrast. (**F**) Spot assay of cells grown under indicated conditions.

Because Pol III transcribes target genes like 5S rRNA and tRNAs that play roles in translation (Turowski and Tollervey, 2016), we wondered whether Pol III perturbation blocks ROLD-induced Hsf1 activation by perturbing CAT tail synthesis. Pol III perturbation could block CAT tail synthesis by preventing ribosome stalling, a necessary prerequisite for the RQC pathway to initiate. We evaluated this possibility using a quantitative stalling reporter similar to one previously developed (Juszkiewicz and Hegde, 2017; Sundaramoorthy et al., 2017) (Figure 3-figure supplement 1C). *BUD27* deletion did not alleviate stalling induced by (CGN)12 (the stalling sequence in RQCsub) in wt or *ROLD* strains relative to control (Figure 3-figure supplement 1C). Thus, the effects of Pol III perturbation on translation should not confound our analysis of RQCsub by preventing ribosomes translating RQCsub from stalling. When we monitored RQCsub mobility by SDS-PAGE to directly observe any changes in CAT tails, we observed that RQCsub expressed in the *ROLD bud27*Δ strain migrated as a higher molecular weight smear compared to the CATylation-deficient *ltn1*Δ *rqc2*Δ strain and similar to the CATylation-competent *ltn1*Δ strain (Figure 3B). Thus, Pol III perturbation leaves CATylation intact.

Given its ability to mitigate Hsf1 hyperactivation in the *ROLD* strain (Figure 3A;Figure 3-figure supplement 1A), we investigated whether Pol III perturbation could inhibit CAT tail aggregation similarly to GuHCl. *ROLD* cells carrying hypomorphic *BUD27* or *RPC17* alleles formed RQCsub inclusions less frequently (3.57% and 4.37% relative to 56.9% in *ROLD*), consistent with a reduction in CAT tail aggregation (Figure 3C). Additionally, *BUD27* deletion in the *ROLD* background increased the mobility of RQCsub by SDS-PAGE, reducing the proportion that remained in the well and increasing the proportion that migrated into the resolving gel (Figure 3-figure supplement 1D). Pol III perturbation thus serves as another tool, in addition to GuHCl, to inhibit CAT tail aggregation.

Given that GuHCl solubilized CAT tails and rescued their degradation in *ROLD* cells (Figure 1C,G), we asked whether Pol III perturbation yields the same effect. To quantify CAT tail degradation, we measured RQCsub_LONG_ stability in *ROLD* and *ROLD bud27*Δ cells (Figure 3D). The difference in RQCsub_LONG_ stability (GFP:RFP) in each of these backgrounds with Hul5 (degradation intact) and without Hul5 (degradation blocked) defined the degree of degradation (Sitron and Brandman, 2019). We validated this approach by demonstrating that the proteasome inhibitor bortezomib did not significantly stabilize RQCsub_LONG_ in *ROLD hul5*Δ and *ROLD bud27*Δ *hul5*Δ strains (Figure 3-figure supplement 1E), indicating that *HUL5* deletion blocked degradation in the *ROLD* and *ROLD bud27*Δ backgrounds. *HUL5* deletion stabilized RQCsub_LONG_ in *ROLD bud27*Δ more than in *ROLD* cells (33.7% compared to 13.6%; Figure 3E). In agreement with the effect of GuHCl (Figure 1G), these data demonstrate that limiting CAT tail aggregation by perturbing Pol III restores degradation of CATylated proteins.

We then used Pol III perturbation to analyze how reducing CAT tail aggregation affects proteostasis by determining its effects on growth in the presence of stress (Figure 3D). As previously published, *BUD27* deletion slightly lowered growth rate at 30°C and strongly impaired growth at 37°C in the wt background (Figure 3F) (Deplazes et al., 2009). We observed similar effects of *BUD27* deletion in the *ROLD* background (Figure 3F). Thus, the sensitivity to mild thermal stress caused by Pol III impairment renders this perturbation unable to rescue heat sensitivity in the *ROLD* background. Despite causing a growth defect in wt cells, *BUD27* deletion enabled faster growth of *ROLD* cells in the presence of cycloheximide (Figure 3F). Unexpectedly, the growth rate of *ROLD bud27*Δ cells upon cycloheximide exposure was greater than that of the *bud27*Δ strain (Figure 3F). Pol III perturbation, similarly to GuHCl, thereby reduces cells’ sensitivity to increased ribosome stalling in addition to inhibiting CAT tail aggregation. These results suggest that improving CAT tail solubility can enhance adaptation to ribosome stall-inducing stress.

## Discussion

We have examined how CAT tail aggregation affects the cell’s ability to cope when its burden of incomplete proteins exceeds Ltn1’s ability to ubiquitylate them. In cells with compromised Ltn1 function, we modulated the aggregation propensity of CAT tails to determine how aggregation affects the cellular capacity to handle elevated levels of incomplete proteins. Aggregation compromised degradation of CATylated proteins (Figure 1F) and diminished cellular fitness during stress (Figure 2E), while inhibiting aggregation reversed each of these effects (Figure 1G;Figure 2E;Figure 3E,F). Our findings suggest that CAT tail aggregation is detrimental, rather than adaptive, in cells with diminished Ltn1 function.

Since their discovery, multiple functions have been attributed to CAT tails. These include a role in assisting Ltn1 activity (Kostova et al., 2017; Osuna et al., 2017; Sitron and Brandman, 2019) and Ltn1-independent roles such as targeting Ltn1-evading RQC substrates for degradation (Sitron and Brandman, 2019) or serving as aggregation-inducing post-translational modifications (Yonashiro et al., 2016). We previously presented evidence supporting an adaptive, Ltn1-independent function for CAT tails by showing that loss of CAT tails sensitized cells to increased ribosome stalling (cycloheximide) specifically in the absence of Ltn1 (Sitron and Brandman, 2019). Here we show that this adaptive Ltn1-independent role is unlikely to be CAT tail-mediated aggregation, as aggregation instead increased sensitivity to cycloheximide (Figure 2E;Figure 3F). Combined with the recent finding that CAT tail degron function is shared with prokaryotic Ala tails (Lytvynenko et al., 2019), our results suggest that marking escaped RQC substrates for degradation is the primary function of CAT tails.

Our findings elevate the question of why yeast and metazoan CAT tails have retained their threonine content, and thereby aggregation propensity, through evolution. We propose two models to explain this. In the first model, CAT tail aggregation facilitates adaptation to an as-of-yet untested stress with evolutionary relevance. The tools we used here to control aggregation may prove useful in probing this model by systematically assessing the fitness benefit of CAT tail aggregation in diverse conditions. The second model proposes that the inclusion of threonine in CAT tails facilitates a function of CAT tails other than aggregation. For instance, threonine incorporation could produce more potent CAT tail degrons, assist in Ltn1-mediated ubiquitylation, or aid in mounting a protective heat shock response. To preserve threonine content, the fitness benefit from this threonine-mediated CAT tail function would need to outweigh any fitness deficits arising from CAT tail aggregation. Given the recent discovery of Rqc2 homologs that incorporate different amino acids into CAT tails (Lytvynenko et al., 2019; Wu Z, Tantray I, Lim J, Chen S, Li Y, Davis Z, Sitron C, Dong J, Gispert S, Auburger G, Brandman O, Bi X, Snyder M, Lu B., 2019), domain-swapping between these homologs to bias CAT tail amino acid content may assist in determining the function of threonine in CAT tails in future studies. Broadly, understanding how the variable amino acid composition of CAT tails has evolved to meet physiological demands in different organisms may prove a fruitful area of research.

We exploited *RQC2* overexpression and Pol III disruption as tools to genetically modify CAT tail aggregation. While these proved useful in analyzing CAT tails in their aggregated and unaggregated states, the underlying mechanism behind these perturbations is not the focus of this study (nor does it influence our conclusions) and remains unclear. *RQC2* overexpression has been proposed to enhance aggregation because endogenous Rqc2 is limiting for CATylation (Yonashiro et al., 2016). Under such a scenario, *RQC2* overexpression would enable CATylation of more escaped RQC substrates that could then nucleate CAT tail aggregates. Although pleiotropic (Turowski and Tollervey, 2016), Pol III perturbation could function in the opposite manner. By limiting translation initiation (Deplazes et al., 2009), Pol III perturbation could decrease the total amount of CATylated proteins in the cell and thereby disfavor nucleation of CAT tail aggregation. Further studies will be needed to decipher the mechanisms by which these perturbations alleviate CAT tail aggregation.

We report here that CAT tail aggregation modifies the toxicity associated with disrupted Ltn1 function. In mice, a hypomorphic *LTN1* allele induces a progressive neurodegeneration that shares phenotypic similarities with amyotrophic lateral sclerosis (ALS) (Chu et al., 2009). It is possible that degenerating neurons in these *LTN1*-hypomorphic mice harbor toxic CAT tail aggregates, as has been observed in a *Drosophila* model of Parkinson Disease (Wu Z, Tantray I, Lim J, Chen S, Li Y, Davis Z, Sitron C, Dong J, Gispert S, Auburger G, Brandman O, Bi X, Snyder M, Lu B., 2019). As the connection between the phenotypes in these disease models and human disease becomes more clear, future studies can focus on strategies to mitigate toxicity associated with compromised Ltn1. Our study serves as a proof-of-principle that disrupting CAT tail aggregation lessens this toxicity.

## Materials and Methods

### Yeast Strains and Culturing

The parental wild-type yeast strain used in this study was BY4741. All gene deletions, genomic integrations, and plasmid transformations were done in this background using standard methods. For a complete list of strains used, see Supplementary Table 1.

Yeast cultures were grown at 30°C in YPD or synthetic defined (SD) media with appropriate nutrient dropouts. SD media used for growth of yeast cultures contained: 2% w/v dextrose (Thermo Fisher Scientific, Waltham, MA), 13.4 g/L Yeast Nitrogen Base without Amino Acids (BD Biosciences, San Jose, CA), 0.03 g/L L-isoleucine (Sigma-Aldrich, St. Louis, MO), 0.15 g/L L-valine (Sigma-Aldrich), 0.04 g/L adenine hemisulfate (Sigma-Aldrich), 0.02 g/L L-arginine (Sigma-Aldrich), 0.03 g/L L-lysine (Sigma-Aldrich), 0.05 g/L L-phenylalanine (Sigma-Aldrich), 0.2 g/L L-Threonine (Sigma-Aldrich), 0.03 g/L L-tyrosine (Sigma-Aldrich), 0.018 g/L L-histidine (Sigma-Aldrich), 0.09 g/L L-leucine (Sigma-Aldrich), 0.018 g/L L-methionine (Sigma-Aldrich), 0.036 g/L L-tryptophan (Sigma-Aldrich), and 0.018 g/L uracil (Sigma-Aldrich). Media additionally contained 5 mM or 10 mM guanidinium hydrochloride (Sigma-Aldrich) where indicated and bortezomib (LC Laboratories, Woburn, MA) treatment lasted 4 hours.

### Plasmids

Plasmids were constructed using NEBuilder HiFi DNA Assembly Master Mix (New England Biolabs, Ipswich, MA). All plasmids used in this study are described in Supplementary Table 2.

Two variants of RQCsub were used that had identical construction except for the region following the stall: one had a fluorescent protein (RFP) and the other had a series of affinity tags. The version without the post-stall RFP was used for microscopy and the version with the post-stall RFP was used in immunoblots; the two exceptions to this are that the version without the post-stall RFP was used in immunoblots in Figure 1C and Figure 1-figure supplement 1B.

### Flow cytometry

Fluorescence was measured using a BD Accuri C6 flow cytometer (BD Biosciences). Three independent yeast cultures were grown to log phase overnight in appropriate SD dropout media. Data analysis was performed with MATLAB (MathWorks, Natick, MA).

For measurements of the Hsf1 activity reporter, raw fluorescence values were first normalized to side scatter, log_2_-transformed, and then the value obtained from the wt strain was subtracted from each condition. These calculations are reported as log_2_ fold wt in the figures.

For measurements of RQCsub_LONG_, a detailed description of the analysis workflow can be found in (Sitron and Brandman, 2019). Briefly, yeast expressing the plasmid were gated according to above-background RFP fluorescence. Then, bleedthrough from the RFP channel was calculated using an RFP-only control strain and subtracted from signal in the GFP channel. The resultant bleedthrough-corrected GFP:RFP values were normalized to the corresponding untreated *hul5*Δ strain for each background. To quantify the change in GFP:RFP upon *HUL5* deletion, the normalized value from the *HUL5* condition was subtracted from that of the *hul5*Δ condition.

For the stalling reporter, the analysis workflow matched that of RQCsub_LONG_ with the order of the fluorophores being reversed, i.e. plasmid-expressing yeast selected based on GFP, bleedthrough from GFP subtracted from RFP. Where indicated, corrected RFP:GFP values were normalized to a non-stalling reporter that contained a non-stalling sequence encoding serine and threonine (Sitron et al., 2017) in place of the stalling CGN codons.

### Statistical Analysis

A paired t-test (using the “ttest” function in MATLAB) was used to assess whether mean measurements differed between two different conditions, using the null hypothesis: μ_condition 1_ = μ_condition 2_. To calculate the s.e.m. for Δ stability measurements (Fig. 1F,G;Fig. 3E), a propagation of error formula was used: SEM_*hul5*Δ - *HUL5*_ = sqrt(SEM_*HUL5*_^2^ + SEM_*hul5*Δ_^2^). To analyze the significance of Δ stability measurements, a t-test for particular contrast was performed using the “lm” and “linearHypothesis” functions in R (R Foundation for Statistical Computing). The null hypothesis for this test was: μ_*hul5*Δ+ condition 1_ – μ_*HUL5* + condition 1_ = μ_*hul5*Δ+ condition 2_ – μ_*HUL5* + condition 2_.

### Immunoblots

For whole-cell immunoblots, 0.375/OD600 x mL of yeast culture (OD600 = 0.4-0.8) grown overnight were pelleted and resuspended then boiled for 5 min at 95°C in 15 μL 4x NuPage LDS Sample Buffer (Thermo Fisher Scientific) with 5% β-mercaptoethanol.

For SDS-PAGE, samples were loaded into a NuPAGE Novex 4-12% Bis-Tris 1.5 mm protein gel (Thermo Fisher Scientific) and run in MOPS buffer. Gels were semi-dry transferred to 0.45 μm nitrocellulose membranes (Thermo Fisher Scientific) using the Trans-Blot Turbo system (Bio-Rad, Hercules, CA). Gels were wet transferred to .45 μm nitrocellulose membranes (Thermo Fisher Scientific) in the Trans-Blot Cell (Bio-Rad) in Towbin Buffer or semi-dry transferred using a Trans-Blot Turbo (Bio-Rad).

Membranes were blocked in TBST with 5% fat-free milk (Safeway, Pleasanton, CA) for 1 hour, incubated overnight at 4°C or at room temperature for 4 hrs in one of the following primary antibodies: 1:3000 mouse α-GFP (MA5-15256, Thermo Fisher Scientific), 1:3000 rabbit α-hexokinase (H2035-01, US Biological, Salem, MA), or 1:1000 rabbit α-RFP (AB233, Evrogen, Moscow, Russia). The following secondary antibodies were then used at 1:5000 dilution: IRDye 800CW donkey anti-mouse, IRDye 800CW goat anti-rabbit, IRDye 680RD goat anti-rabbit, or IRDye 680RD goat anti-mouse (LiCor Biosciences, Lincoln, NE). Blots were scanned on a Licor Odyssey (LiCor Biosciences).

### Lysate Preparation and Immunoprecipitation (IP)

Yeast were grown in SD media and harvested at OD600 0.8 to 1.0. Cells were harvested using either 1) centrifugation followed by resuspension in IP buffer (100 mM KOAc, 10 mM MgCl2, 25 mM HEPES-KOH pH 7.4) and freezing of cell droplets in liquid nitrogen; or 2) vacuum filtration and flash freezing in liquid nitrogen. Frozen yeast were cryogenically pulverized into powder using a Freezer/Mill (SPEX SamplePrep, Metuchen, NJ) and stored at −80°C.

Frozen yeast powder was thawed and solubilized at 4°C in IP buffer supplemented with Pierce Protease Inhibitor Tablets, EDTA-free (Thermo Fisher Scientific). Crude lysate was clarified by a single spin at 5000 x g, incubated with GFP-Trap_A (ChromoTek, Planegg-Martinsried, Germany) resin for 1 hour at 4°C with rotation, then washed 10 times in IP buffer. For immunoblotting, washed GFP-Trap resin was boiled in 4X NuPAGE LDS Sample Buffer (Thermo Fisher Scientific) with 5% β-mercaptoethanol and analyzed as described above in “Immunoblots.”

### TEV Protease Digestion

GFP IP was performed as described above, but instead of immediate elution, the rein was equilibrated in IP buffer with 1 mM DTT and incubated overnight at 4°C with 5 μL ProTEV Plus (Promega, Sunnyvale, CA) in 100 μL total reaction volume. The resin was then washed with fresh IP buffer with 1 mM DTT and boiled in 4X NuPAGE LDS Sample Buffer (Thermo Fisher Scientific) with 5% β-mercaptoethanol and analyzed as described above.

### Fluorescence Microscopy

Live cell fluorescence imaging was performed using two set-ups: 1) a Nikon Eclipse Ti-E inverted fluorescence microscope (Nikon, Melville, NY) using a 100X/1.4NA oil objective lens and μManager (Edelstein et al., 2010), and 2) a Nikon Eclipse 80i microscope using a 100X/1.4NA oil objective lens and Metamorph (Molecular Devices, San Jose, CA). Yeast were grown in SD media to log phase, centrifuged briefly, resuspended in 5 μL fresh SD media and mounted on a clean coverslip and slide. Images were prepared and analyzed with ImageJ (NIH). For quantification, images were randomly acquired until N > 300 cells. Independent cultures were imaged on different days and error represents standard deviation of 3 independent cultures.

### Yeast spot assay

Yeast were grown to log-phase in YPD, then diluted to OD600 = 0.1. 200 μL of this diluted culture was diluted 1:10 into YPD four times in a sterile 96-well plate (Greiner Bio-one, Kremsmünster, Austria). 5 μL of the serial dilutions were transferred from the 96-well plate onto YPD agar plates with or without additional 5 mM guanidinium hydrochloride (Sigma-Aldrich) and/or 50 ng/mL cycloheximide (Sigma-Aldrich). Plates were then incubated at 30°C or 37°C. Plates grown at 30°C with or without 5 mM guanidinium hydrochloride as well as 37°C plates were imaged after two days. Plates containing 50 ng/mL cycloheximide with or without 5 mM guanidinium hydrochloride grown at 30°C were imaged after three days. Plates with 5 mM guanidinium hydrochloride grown at 37°C were imaged after six days.

## Acknowledgements

We thank D. Pincus, D. Jarosz, R. Kopito, B. Lu, Z. Wu, E.P. Geiduschek, and the members of the Brandman lab for helpful suggestions. We thank the Theriot and Straight labs (Stanford University) for the use of their microscopes. We thank J. Weissman and D. Wong for facilitating the screen for suppressors of RQC-mediated Hsf1 activation. This work was supported by Stanford University (O.B.), the US National Institutes of Health (grant No. R01GM115968 to O.B.), and the National Institute of General Medical Sciences of the US National Institutes of Health (grant No. T32GM007276 to C.S.S.).

## Author Contributions

O.B., C.S.S., and J.P. conceived of the study and designed the experiments. C.S.S., J.P., and J.M.G. performed the experiments. O.B., C.S.S., and J.H.P. analyzed and interpreted the data. O.B., C.S.S., and J.H.P. wrote the manuscript. O.B. supervised the study.

---

**Figure 1-figure supplement 1.**
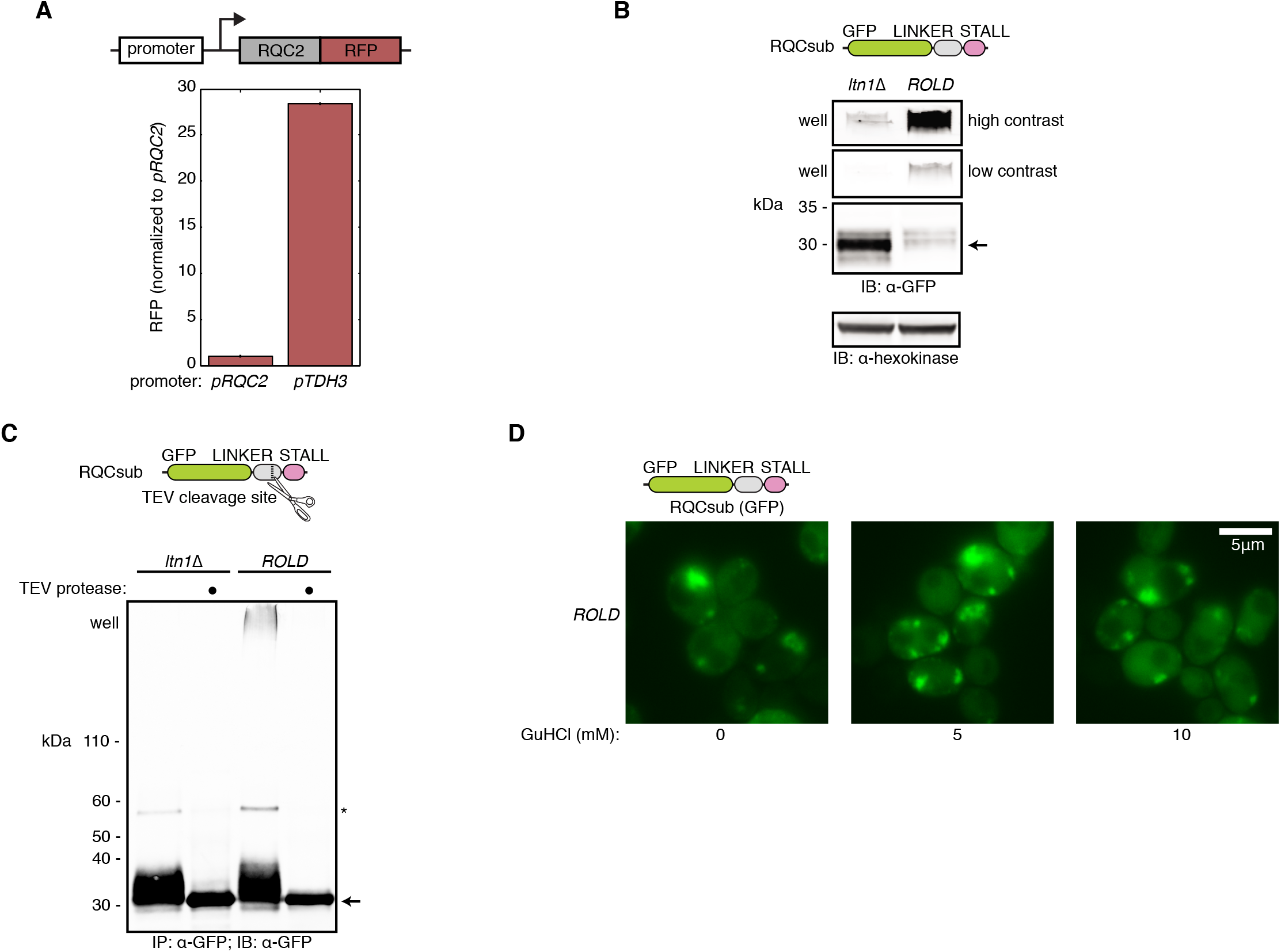
*RQC2* overexpression and guanidinium hydrochloride are tools to control CAT tail aggregation. (**A**) Flow cytometry of cells expressing Rqc2-RFP to assess the degree of Rqc2 overproduction by the *TDH3* promoter. Error bars represent s.e.m. from three independent cultures. (**B**) Immunoblot (IB) of RQCsub expressed in *ltn1*Δ compared to *ROLD*. (**C**) IB of RQCsub immunoprecipitated (IPed) from *ltn1*Δ and *ROLD* lysates with and without tobacco etch virus (TEV) protease treatment. Arrow denotes molecular weight of non-CATylated RQCsub. Asterisk indicates the full-length RQCsub protein product, produced when ribosomes translate through the stall sequence (region past the stall not pictured in schematic). (**D**) Microscopy of RQCsub expressed in *ROLD* grown in various concentrations of guanidinium hydrochloride (GuHCl).

**Figure 1-figure supplement 2.**
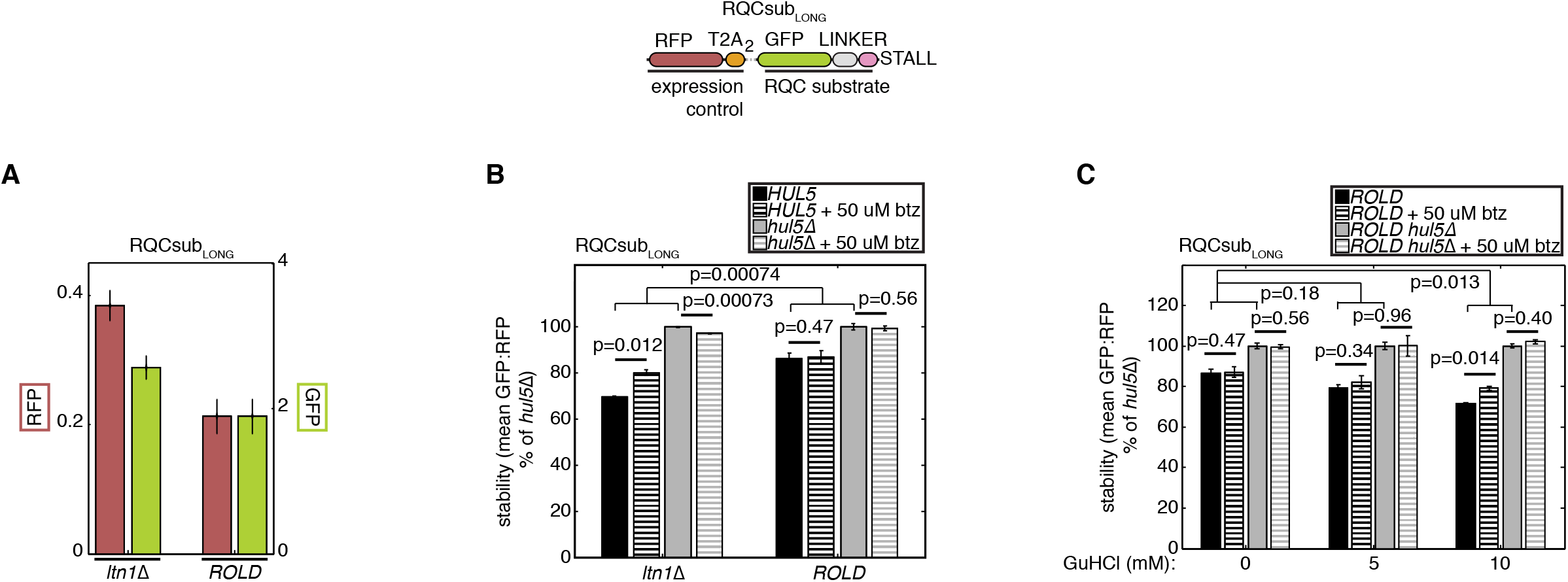
Supporting data for the effect of aggregation of CAT tail-mediated degradation. (**A**) Flow cytometry of cells expressing RQCsub_LONG_. Error bars indicate s.e.m. from three independent cultures. (**B**) and (**C**) Additional data to support Figure 1F,G. Stability measurements of RQCsub_LONG_ expressed in *ROLD* cells grown in indicated GuHCl concentrations with additional bortezomib treatment to inhibit the proteasome and *HUL5* deletion to measure CAT tail-mediated degradation. Error bars as in **A**. P-values are indicated above bars. Thick lines indicate paired t-tests, probing the significance of bortezomib (btz)-induced stabilization. Thin lines denote t-tests for particular contrast, measuring how significantly different *HUL5* deletion-induced stabilization is under different conditions.

**Figure 3-figure supplement 1.**
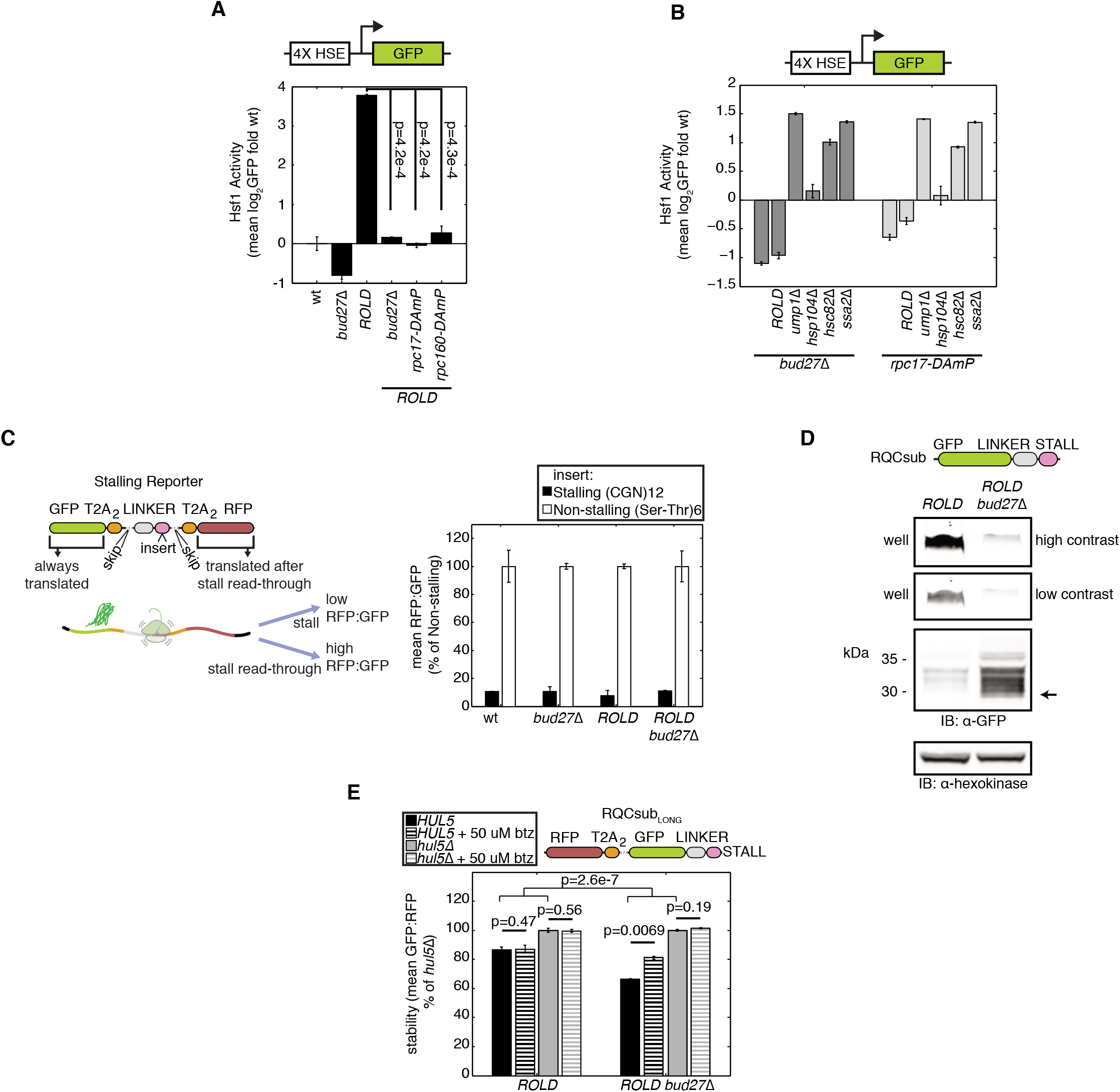
Effects of Pol III perturbation on Hsf1 activation, stalling, and CAT tail degron activity. (**A**) Flow cytometry of cells containing an integrated reporter for Hsf1 activation. Error bars indicate s.e.m. from three independent cultures. P-values from paired t-tests indicated above bars. (**B**) Flow cytometry of Pol III-perturbed cells containing an integrated reporter for Hsf1 activation. These data are also contained in Figure 3A, but are reordered here to simplify comparisons within two Pol III-perturbed genetic backgrounds. Error bars as in **A**. (**C**) Left, schematic of a stalling reporter similar to that used in Juszkiewicz and Hegde 2017 and Sundaramoorthy et al. 2017. Right, schematic of stalling reporter with the same (CGN)12 stalling sequence contained in RQCsub or a non-stalling (Ser-Thr)6 sequence. Error bars as in **A**. (**D**) IB of RQCsub expressed in *ROLD* compared to *ROLD bud27*Δ. (**E**) Additional data to support Figure 3E. Stability measurements of RQCsub_LONG_ expressed in indicated strains with bortezomib (btz) treatment to inhibit the proteasome and *HUL5* deletion to block CAT tail-mediated degradation. Error bars indicate s.e.m. from three independent cultures. P-values are given above bars. Results of paired t-tests measuring the significance of bortezomib-induced stabilization are indicated with thick lines. The result of a t-test for particular contrast is indicated with thin lines; this assesses how significantly different *HUL5* deletion-induced stabilization is in *ROLD* compared to *ROLD bud27*Δ.

**Supplementary Table 1.** Yeast strains used in this study.

| Strain Number | Description |
| --- | --- |
| yOB199 | BY4741 |
| yOB259 | BY4741 *ltn1Δ::kanMX* |
| yOB454 | BY4741 *rqc2Δ::kanMX* |
| yOBj004 | BY4741 *ltn1Δ::kanMX rqc2Δ::natMX* |
| yOB518 | BY4741 *pRQC2::HIS3-pTDH3* |
| yOB551 | BY4741 *pRQC2::HIS3-pTDH3 ltn1Δ::natMX* |
| yOB255 | BY4741 *ura3::HSE-EmGFP* |
| yOB549 | BY4741 *ura3::HSE-EmGFP rqc2Δ::kanMX* |
| yOB550 | BY4741 *ura3::HSE-EmGFP ltn1Δ::kanMX* |
| yOBj245 | BY4741 *ura3::HSE-EmGFP rqc2Δ::kanMX ltn1Δ::natMX* |
| yOB515 | BY4741 *ura3::HSE-EmGFP pRQC2::HIS3-pTDH3* |
| yOBj185 | BY4741 *ura3::HSE-EmGFP pRQC2::HIS3-pTDH3 ltn1Δ::natMX* |
| yOB1118 | BY4741 *RQC2-WT::LEU ltn1Δ::hphMX* |
| yOB2825 | BY4741 *RQC2-WT::LEU ltn1Δ::hphMX hul5Δ::natMX* |
| yOB3812 | BY4741 *pRQC2::HIS3-pTDH3 ltn1Δ::natMX hul5Δ::hphMX* |
| yOBj482 | BY4741 *ura3::HSE-EmGFP ump1Δ::natMX* |
| yOBj538 | BY4741 *ura3::HSE-EmGFP ump1Δ::natMX bud27Δ::kanMX* |
| yOB738 | BY4741 *ura3::HSE-EmGFP ump1Δ::natMX rpc17-DAmP::natMX* |
| yOB736 | BY4741 *ura3::HSE-EmGFP hsp104Δ::natMX* |
| yOB808 | BY4741 *ura3::HSE-EmGFP hsp104Δ::natMX bud27Δ::kanMX* |
| yOB809 | BY4741 *ura3::HSE-EmGFP hsp104Δ::natMX rpc17-DAmP::kanMX* |
| yOBj542 | BY4741 *ura3::HSE-EmGFP hsc82Δ::natMX* |
| yOB818 | BY4741 *ura3::HSE-EmGFP hsc82Δ::natMX bud27Δ::kanMX* |
| yOB722 | BY4741 *ura3::HSE-EmGFP hsc82Δ::natMX rpc17-DAmP::kanMX* |
| yOBj563 | BY4741 *ura3::HSE-EmGFP ssa2Δ::natMX* |
| yOB819 | BY4741 *ura3::HSE-EmGFP ssa2Δ::natMX bud27Δ::kanMX* |
| yOB725 | BY4741 *ura3::HSE-EmGFP ssa2Δ::natMX rpc17-DAmP::kanMX* |
| yOBj536 | BY4741 *ura3::HSE-EmGFP bud27Δ::kanMX* |
| yOB710 | BY4741 *ura3::HSE-EmGFP rpc17-DAmP::kanMX* |
| yOBj495 | BY4741 *pRQC2::HIS3-pTDH3 ltn1Δ::natMX bud27Δ::kanMX* |
| yOBj556 | BY4741 *pRQC2::HIS3-pTDH3 ltn1Δ::natMX rpc17-DAmP::kanMX* |
| yOBj536 | BY4741 *ura3::HSE-EmGFP bud27Δ::kanMX* |
| yOBj185 | BY4741 *ura3::HSE-EmGFP pRQC2::HIS3-pTDH3 ltn1Δ::natMX bud27Δ::kanMX* |
| yOBj556 | BY4741 *ura3::HSE-EmGFP pRQC2::HIS3-pTDH3 ltn1Δ::natMX rpc17-DAmP::kanMX* |
| yOBj626 | BY4741 *ura3::HSE-EmGFP pRQC2::HIS3-pTDH3 ltn1Δ::natMX rpc160-DAmP::kanMX* |
| yOB3812 | BY4741 *pRQC2::HIS3-pTDH3 ltn1Δ::natMX hul5Δ::hphMX bud27Δ::kanMX* |

**Supplementary Table 2.**
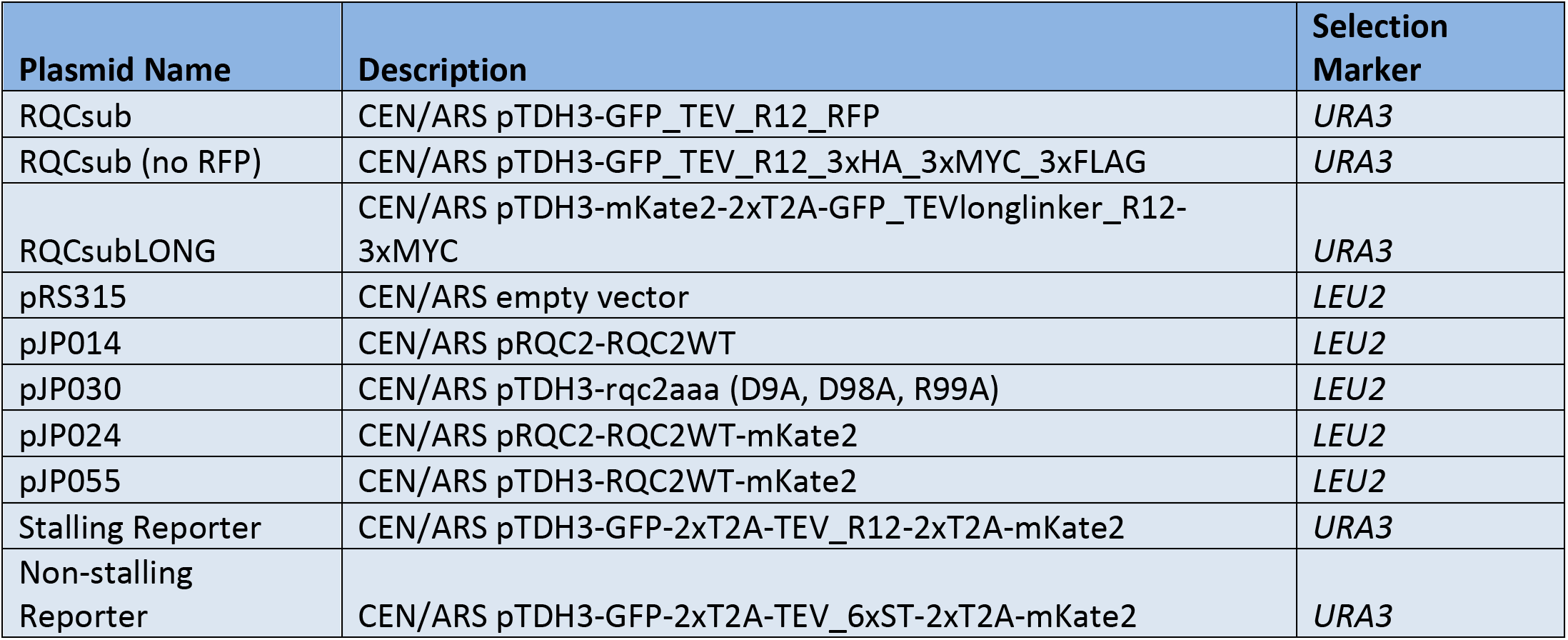
Plasmids used in this study.

| Plasmid Name | Description | Selection Marker |
| --- | --- | --- |
| RQCsub | CEN/ARS pTDH3-GFP_TEV_R12_RFP | *URA3* |
| RQCsub (no RFP) | CEN/ARS pTDH3-GFP_TEV_R12_3xHA_3xMYC_3xFLAG | *URA3* |
| RQCsubLONG | CEN/ARS pTDH3-mKate2-2xT2A-GFP_TEVlonglinker_R12-3xMYC | *URA3* |
| pRS315 | CEN/ARS empty vector | *LEU2* |
| pJP014 | CEN/ARS pRQC2-RQC2WT | *LEU2* |
| pJP030 | CEN/ARS pTDH3-rqc2aaa (D9A, D98A, R99A) | *LEU2* |
| pJP024 | CEN/ARS pRQC2-RQC2WT-mKate2 | *LEU2* |
| pJP055 | CEN/ARS pTDH3-RQC2WT-mKate2 | *LEU2* |
| Stalling Reporter | CEN/ARS pTDH3-GFP-2xT2A-TEV_R12-2xT2A-mKate2 | *URA3* |
| Non-stalling Reporter | CEN/ARS pTDH3-GFP-2xT2A-TEV_6xST-2xT2A-mKate2 | *URA3* |

